# Associations between neonatal cry acoustics and visual attention during the first year

**DOI:** 10.1101/658732

**Authors:** Aicha Kivinummi, Gaurav Naithani, Outi Tammela, Tuomas Virtanen, Enni Kurkela, Miia Alhainen, Dana J. H. Niehaus, Anusha Lachman, Jukka M. Leppänen, Mikko J. Peltola

## Abstract

It has been suggested that early cry parameters are connected to later cognitive abilities. The present study is the first to investigate whether the acoustic features of infant cry are associated with cognitive development already during the first year, as measured by oculomotor orienting and attention disengagement. Cry sounds for acoustic analyses (fundamental frequency; F0) were recorded in two neonatal cohorts at the age of 0-5 days (Tampere, Finland) or at 6 weeks (Cape Town, South Africa). Eye tracking was used to measure oculomotor orienting to peripheral visual stimuli and attention disengagement from central stimuli at 8 months (Tampere) or at 6 months (Cape Town) of age. In the Tampere cohort, a marginal positive correlation between fundamental frequency of cry (F0) and visual attention disengagement was observed; infants with a higher neonatal F0 were slower to shift gaze away from the central stimulus to the peripheral stimulus. However, a similar correlation was not observed in the Cape Town cohort. No associations between F0 and oculomotor orienting were observed in either cohort. We discuss possible factors influencing the discrepancy in results between the cohorts and suggest directions for future research investigating the potential of early cry analysis in predicting later cognitive development.

## Introduction

Monitoring children’s cognitive development is an important part of pediatric practice. Traditional psychometric methods enable detailed and reliable measurements especially after two years of age. However, the assessment of cognitive development during the neonatal period and early infancy is faced with multiple challenges, such as age-related changes in how different cognitive functions can be measured. For example, motor abilities are emphasized in most early measures, but their relation to cognitive functioning may vary at different ages, which may weaken the predictive validity of early assessments of cognitive development (Colombo, 1993; McCall, 1979). These challenges make research on the early markers of later cognitive functioning important.

The current study is focused on the connection between the acoustic characteristics of infant cry and visual attention within the first year of life. Cry is a vital, reflex-like means of communication present from the very beginning of life (Newman, 2007) and it indicates the neurological status of the neonate (LaGasse et al., 2005; Lester, 1987). Variations in visual attention during the first year have been shown to be associated with later cognitive development (Colombo, 2002; Dougherty & Haith, 1997; Rose et al., 2003; Rose et al., 2012), thus providing an outcome measure that emerges early in development and is meaningfully associated with more complex and later-emerging cognitive functions. As both the acoustic characteristics of cry and visual attention can be measured reliably at an early age, examining their interrelations may help to determine the potential of neonatal cry analysis as an accessible, complementary marker of later development.

A limited number of small-scale studies are available to suggest that early cry parameters are associated with later cognitive abilities. Lester (1987) observed that preterm and term infants having high average fundamental frequency of cry (F0; heard as the pitch of the cry) and larger variability in F0 within 2 weeks of term postconceptional age were more likely located in the group with lower scores on the Bayley Scales of Infant Development at 18 months of age compared to infants with lower average F0 and less variable F0. In addition, high variation in neonatal F0 predicted lower scores in the McCarthy Scales of Children’s Abilities at 5 years of age. Similarly, studying infants prenatally exposed to methadone, Huntington et al. (1990) found that higher and more variable F0 in the first days of life were related to lower mental development scores measured by the Bayley scales at 8, 12, and 18 months. However, the association was only marginal at 24 months. Both Lester (1987) and Huntington et al. (1990) observed that the relation between the F0 variables and the cognitive development outcomes remained after controlling for risk factors including prematurity and prenatal exposure to drugs. Other studies have observed that the acoustic characteristics of cry in children with neurodevelopmental disorders such as CNS abnormalities and developmental delays (for reviews, see (LaGasse et al., 2005; Michelsson & Michelsson, 1999) and children with autism spectrum disorders (Esposito et al., 2014; Sheinkopf et al., 2012) are associated with an atypical and more unstable F0. The similarities in the findings of studies investigating at-risk infants, as well as the scarce findings among normally developing infants emphasize the importance of further research on the connections between the acoustic features of newborn cry and cognitive functioning during the first year of life among typically developing infants.

Regarding the neural basis of variation in neonatal cry, it is suggested that imperfect functioning of the vagus nerve and inhibitory parasympathetic nervous system activity are reflected in infant cry as higher and more variable F0 (Porter et al., 1988; Lester et al., 1976). Neonatal cry requires versatile integration of respiratory and larynx area organs, which are suggested to be controlled in early, reflex-like crying exclusively by the brainstem. No higher areas than the midbrain are required for producing cry during the first months of life (Newman, 2007).

Consequently, the integrity of the brainstem is suggested to influence the acoustic features of cry (Lester, 1987; Vohr et al., 1989) and the vagus nerve of the parasympathetic nervous system is considered as a crucial pathway through which the brainstem regulates neonatal and infant cry (Newman, 2007; LaGasse et al., 2005; Lester et al., 1989). The vagus nerve originates from the medulla oblongata in the brainstem and is densely connected with the vocal cords and respiratory muscles, on which it has an inhibitory function. In addition, neonatal F0 has been shown to vary as a function of cardiac vagal tone (Shinya et al., 2016). Thus, imperfect inhibitory parasympathetic control of the cry organs is likely reflected in higher and more variable infant F0. These findings highlight the role of the brainstem and vagal system in affecting neonatal cry acoustics.

In addition to influencing infant cry production, integrity of the brainstem may have a pivotal impact on visual attention and the processing of visual information (Colombo, 2001; Geva & Feldman, 2008; Geva et al., 2014; Hunnius & Geuze, 2004). The physiologically based model of attention suggests that vagus nerve mediated organizational processes of the brainstem are required to integrate sensory and motor information on attention. As a part of the parasympathetic nervous system, the vagus nerve has an inhibitory function on arousal. Further on, attention is typically modulated by arousal state and it has been observed that infants with brainstem dysfunction show atypical modulation of attention by arousal. Healthy neonates preferred familiar visual stimuli during increased arousal and novel stimuli during decreased arousal, whereas this pattern was less pronounced in neonates with brainstem dysfunction (Gardner et al., 2003). Karmel et al. (1996) observed that a similar atypical arousal-modulated attention pattern persists at least until 4 months of age in infants with brainstem dysfunction. In addition, sensitivity to social and non-social content of visual stimuli is connected to brainstem integrity. Geva et al. (2017) detected that children with discrete neonatal brainstem dysfunction preferred an active nonsocial content (a flock of birds flying) or a neutral social content (a person listening) over an active social content (a person expressing affective narratives). These findings were opposite to the pattern of attention in normal controls and the difference remained even in a follow up at 8 years of age.

Thus, it appears that neonatal cry and infant visual attention share partially overlapping neural mechanisms, with particularly the brainstem implicated in both. In the current study, we therefore investigated the associations between infant cry acoustics (F0 and its variation) during the neonatal period (0-6 weeks) and visual attention at 6-8 months of age. As the indicator of infant visual attention, we measured oculomotor orienting and attention disengagement, both operationalized as saccadic reaction times in separate eye tracking paradigms.

The ability to produce rapid shifts of gaze to stimuli appearing in a new spatial location, typically measured as saccadic reaction time, matures in early infancy and its variations correlate with later cognitive development (Dougherty & Haith, 1997). For example, infants with longer latency of saccades to peripheral target stimuli were shown to score lower on standardized IQ measures at 4 years of age (Dougherty & Haith, 1997). In addition, saccadic reaction times have been shown to be affected by neurodevelopmental risk factors such as prenatal alcohol exposure (Green et al., 2013), preterm birth (Hunnius et al., 2008) and familial risk for autism spectrum disorder (Elsabbagh et al., 2013). Hence, studying infant visual attention with paradigms measuring saccadic reaction times may provide objective and applicable markers for examining neurocognitive development in a variety of populations.

We used two infant attention paradigms for measuring saccadic reaction times. First, oculomotor orienting was measured as saccadic reaction time to a new peripheral stimulus following the offset of a stimulus on the center of the screen. Second, we measured disengagement of attention as the duration of gaze shift from centrally presented face and non-face stimuli to laterally presented competing stimuli. Particularly oculomotor orienting has been associated with later cognitive development, with faster saccadic reaction times in infancy predicting higher childhood IQ (Dougherty & Haith, 1997; Rose et al., 2012). Attention disengagement paradigms have been especially used for investigating infants’ attention to social stimuli such as faces (Peltola et al., 2008) with recent findings indicating that variations in infants’ attention to faces in this paradigm are associated with later social development (Peltola et al., 2018). Furthermore, given that oculomotor orienting and attention disengagement appear to mature at different speed during the first year (with attention disengagement maturing later) (Colombo, 2001; Hood & Atkinson, 1993; Matsuzawa & Shimojo, 1997) and rely on different neural mechanisms (Colombo, 2001; Rafal & Robertson, 1995) we considered it important to include both measures of infant attention in the current study.

Taken together, as infant cry and visual attention may have a partially overlapping neural basis and as the basic functions of infant visual attention may be important in understanding later-emerging more complex cognitive and social abilities, infant visual attention is an ideal outcome measure for attempts to determine whether the acoustic features of newborn cry are associated with normative developmental variations in basic cognitive functions.

Due to the rather explorative nature of our study, we implemented the analysis across two independent samples of infants from Tampere, Finland and Cape Town, South Africa, to assess the replicability of the findings. Based on previous findings showing that both F0 of infant cry (Huntington et al., 1990; Lester, 1987) and saccadic reaction times (Dougherty & Haith, 1997; Rose et al., 2012) are associated with childhood cognitive development, we studied whether higher and more variable F0 would be associated with longer saccadic reaction times in the oculomotor orienting task. Regarding the attention disengagement paradigm, the paucity of previous research prevented testing strong directional hypotheses. However, given the links between neonatal cry production and brainstem integrity on the one hand (Newman, 2007; Vohr et al., 1989) and between brainstem integrity and attention to social stimuli on the other hand (Geva et al., 2017), it could be tentatively hypothesized that higher and more variable F0 would be associated with less attention to faces in the attention disengagement paradigm (i.e., faster saccadic reaction times in shifting attention away from faces).

## Materials and Methods

### Ethics Statement

The study adhered to ethical guidelines and the study protocol was approved by the Ethical Committee of Tampere University Hospital and, on behalf of Cape Town participants, by the Institutional Review Board of Stellenbosch University. Infants were included in the study after a written consent from their parent was obtained.

### Participants

Data for the current analyses were obtained from two cohorts, one in Tampere, Finland, and the other in Cape Town, South Africa. Our interest was to study the associations between acoustic variables of cry during the neonatal period (Tampere: 0-5 days after birth; Cape Town: 6 weeks of age) and infant visual attention measured by eye tracking methods (Tampere: 8 months; Cape Town: 6 months of age).

#### Tampere cohort

Participants in the Tampere cohort were enrolled in the study at birth (0-5 days) and subsequently invited for a follow-up visit at 8 months of age (mean = 253.58 days, *SD* = 12.34 days). A total of 104 neonates were recruited from the Tampere University Hospital Maternity Ward Units and the Neonatal Ward Unit. Infants were considered eligible to be enrolled in the study if the infant did not have any kind of abnormalities in the neck area, nor a history of recent airway suctioning, which may influence the voice, and if one of the parents understood Finnish or English sufficiently to be able to be adequately informed about the study.

From the original pool of 104 infants, 6 infants did not cry when the researcher was present, hence cry recording was captured from 98 infants. Out of these, 73 infants attended the eye tracking follow-up visit at 8 months. Eye tracking data were not obtained from 5 infants because of technical problems and from 4 infants because the infant showed inconvenience about the registration situation. Two infants were further excluded because of not being full-term (≥ 37 weeks), 3 because of low birth weight (≥ 2500 grams), 2 because of perinatal asphyxia, and 1 because of an abnormal brain MRI finding.

From this sample of 56 infants, 2 infants needed to be excluded because of the inclusion criteria of the cry analysis. At least 3 expiratory cry utterances from the beginning of the recording had to be present for analyzing the F0. For the analyses of oculomotor orienting, 2 infants were excluded for having less than 4 trials with good gaze data for calculating oculomotor orienting. For the analyses of attention disengagement, 8 infants were excluded for having less than 3 scorable trials in one of the stimulus conditions (i.e., a non-face control stimulus and happy and fearful face stimuli).

Consequently, 52 infants (22 female, 30 male) and 46 infants (female 19, male 27) were included in the oculomotor orienting and attention disengagement analyses, respectively. The included infants did not have known health or developmental disadvantages nor known prenatal mother-related risk factors, such as drug abuse, untreated diabetes or high blood pressure. Data about health, risk factors, and medical therapies were collected from the medical records of the hospital.

In addition to the primary analyses, we ran additional analyses including only those infants whose cry was captured from the beginning of a cry bout. This was done to enable more direct comparisons with the Cape Town cohort in which the cry recordings were always captured from the beginning of the cry bout. Thus, this analysis controlled the possible influence of the location of the cry sample in the recording. For these analyses, 29 and 28 infants were included in the oculomotor orienting and attention disengagement analyses, respectively.

#### Cape Town cohort

The data for the Cape Town cohort was obtained from a larger research project in collaboration with the Department of Psychiatry at the Stellenbosch University. Participants were enrolled in this study at the age of 6 weeks (mean = 44.73 days, *SD* = 3.34 days) and they participated in the follow-up registration at 6 months (mean = 187.85 days, *SD* = 13.84 days) of age. Similarly to the Tampere cohort, the infants did not have any kind of abnormalities in the neck area, nor a history of recent airway suctioning, which may influence the voice. Likewise, the other exclusion criteria were concordant with the Tampere cohort.

Cry recordings were obtained from a total of 100 infants. The follow-up assessment was completed by 73 infants. Of these, 5 infants needed to be excluded because of the age criteria at cry recording and 4 because there were less than 3 expiratory cry utterances in the recording. Three infants were excluded for not being full-term (≥ 37 weeks). Birthweight data was not available for this cohort, and none of the infants had perinatal asphyxia or a brain MRI finding. The infants did not have other health or developmental disadvantages. Considering the inclusion criteria of mother-related risk factors, we excluded 6 infants due to prenatal alcohol use. For the eye tracking data, identical inclusion criteria were used as in the Tampere cohort. Consequently, 47 infants (18 female, 29 male) and 53 infants (19 female, 34 male) were included in the oculomotor orienting and disengagement analyses, respectively.

Within the cohort in the primary analysis, there were 14 infants exposed to prenatal SSRI medication. Therefore, to control the possible influence of SSRI medication, additional analysis were conducted after excluding these participants, resulting in 37 and 42 infants in the oculomotor orienting and attention disengagement analyses, respectively. In both the analyses a proportion of mothers had psychiatric problems such as depression and anxiety disorders. This was due to the interests in the larger research project, from where the data was captured.

### Measurements

#### Cry recordings in Tampere

We recorded one cry sample from each infant in normal clinical settings at the age of 0 to 5 days. During the recording, infants were laying on supine position. The location of the captured sample in the cry bout varied since we recorded spontaneous cries which were not launched by a distinct painful trigger. Soothing was performed by the parent whenever needed.

The duration of the samples in the whole data ranged from 14.4 s to 430.5 s (mean =114.2 s; *SD* = 81.7) The recording microphone was a Tascam DR-100MK Linear PCM Recorder, Røde M3 cardioid microphone. The samples were stored with a 48-kHz sampling rate, in a 24-bit Waveform Audio file (WAV) format. The microphone was at a stand at 30-cm distance directly in front of the infant’s mouth.

#### Cry recordings in Cape Town

Recording of the cry was done as a part of a standardized protocol at a routine 6-week baby clinic visit to the nurse. The room was always the same and the cry was elicited by vaccination or measuring the infant’s weight on a scale. During the vaccination, the infant was laying on mother’s lap. Soothing was performed in the way the mother preferred. We captured one cry sample at the beginning of the cry bout from every infant.

The duration of the samples in the whole data ranged from 16.9 s to 150 s (mean = 80.7 s; *SD* = 18.8 s). The audio recorder used was a Zoom H4n recorder with built-in condenser microphones. Recordings were stored with a 48-kHz sampling rate, two-channel audio in a 24-bit Waveform Audio file (WAV) format. The distance between the infant’s mouth and the microphone was 1.3 m for infants being vaccinated and 70 cm for infants being weighed. The microphone was fixed on a wall stand.

#### Acoustic variables

As the cry samples were captured in clinical settings containing varying amounts of extraneous sounds, the first task was to separate the cry vocalizations from these extraneous sounds which are not useful for the purpose of acoustic analysis. We used an in-house hidden Markov model (HMM) (Schuster-Böckler & Bateman, 2007) based audio segmentation system (Naithani et al., 2018) for cry segment extraction, and the acoustic analysis was performed in MATLAB (Mathworks, Natick, MA). The cry data was divided to three acoustic regions; expiratory phases, inspiratory phases, and residual, where residual included all the vocalizations that are not expiratory or inspiratory cries. We included only expiratory phases in the analysis, based on the frequent praxis in earlier studies as well as the results of our preliminary study showing that expiratory cries were detected more reliably than inspiratory cries by the analysis tool (Naithani et al., 2018).

In the analyses, we used the first 5 expiratory phases from the beginning of the captured cry sample. The missing values of the used variables were imputed based on other scorable phases if there was a minimum of 3 scorable phases. Only phases with duration longer than 500 ms were included, in line with previous studies (Fuamenya et al., 2015; Reggiannini et al., 2013). The identified expiratory phases were then divided into overlapping frames of 25-ms duration and 50 % overlap, and fundamental frequency (F0) was extracted for each such frame.

As cry variabes we used fundamental frequency (F0) and the variation of fundamental frequency (F0var), which we estimated using the YIN algorithm (De Cheveigné et al., 2002). In line with earlier studies (Michelsson & Michelsson, 1999; Rothgänger, 2003), F0 was defined as the mean of first five within-sample means of frame-wise fundamental frequencies, and F0var was defined as mean of the first five within-sample standard deviations of frame-wise fundamental frequencies. Cry phases with more than 70% inharmonic frames selected via absolute threshold parameter of the YIN algorithm (here 0.3), were excluded (for more information, see Fuamenya et al., 2015). Descriptive characteristics of F0 in different analyses are presented in Table 1.

**Table 1.**
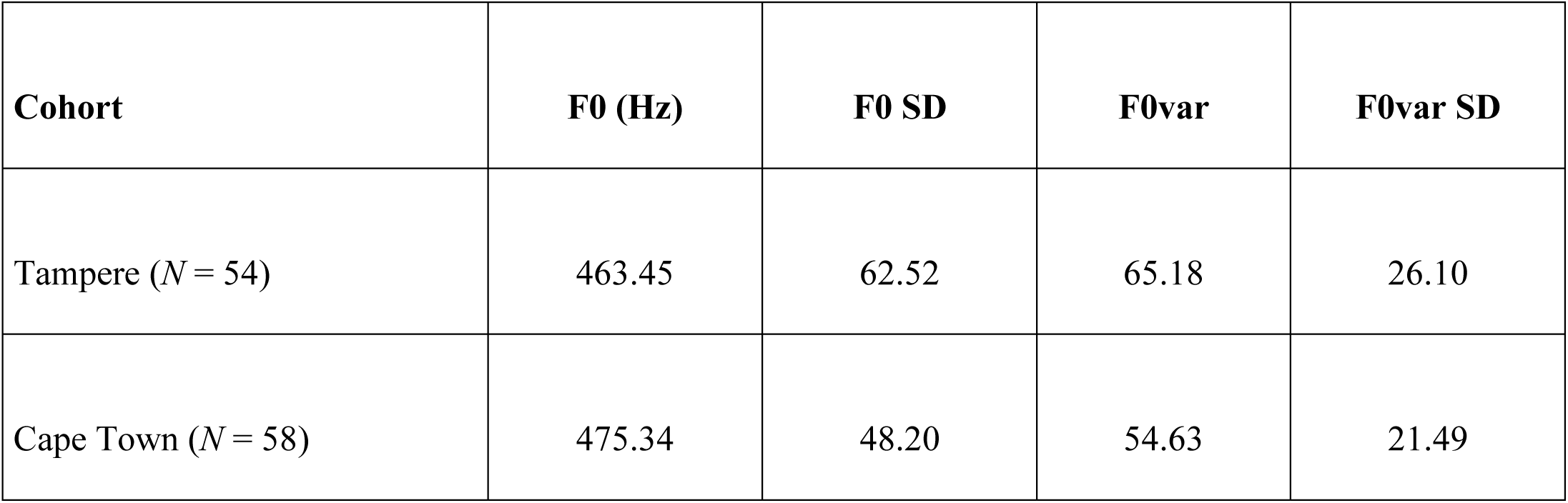
Descriptive data of F0 in Tampere and Cape Town cohorts.

### Eye tracking assessments in Tampere

In the 8-month follow-up visit, the infants participated in an eye tracking assessment of saccadic reaction times measured during separate oculomotor orienting and attention disengagement paradigms. During the assessment, the infant sat on the parent’s lap in a dimly lit room watching the test stimuli on a 23-in monitor located at 60±5 cm distance from the infant’s eyes. The parent was asked to close her/his eyes during the recording to avoid false registrations (Gredebäck et al. 2010). The session was paused or terminated in case the infant expressed inconvenience or was not willing to co-operate. Eye tracking data were collected by using a Tobii TX300 remote eye tracking camera (Tobii Technology, Stockholm, Sweden) with a 300-Hz sampling rate.

Before recording the data, the eye tracker was calibrated by using a 5-point calibration script and Tobii SDK calibration algorithms. We repeated the calibration two times if one or more calibration points were missing or inadequate. We used the final calibration in case none of the calibrations produced an adequate result.

The tasks (i.e., oculomotor orienting and attention disengagement) were both divided in two blocks that were presented in alternate order starting with the first block of the oculomotor orienting task, followed by the first disengagement block, and ending with the second oculomotor orienting and disengagement blocks. Both oculomotor orienting blocks included 16 trials and the disengagement blocks included 24 trials (i.e., a total of 32 and 48 trials in the oculomotor orienting and disengagement tasks, respectively). Cheerful music with smiling children was played between the blocks. Stimulus presentation was controlled by an open-source software written in Python programming language (https://github.com/infant-cognition-tampere/drop), together with Psychopy functions and a Tobii SDK plug-in. For both tasks, all data processing from the raw *x-y* gaze position coordinates to the parameters reflecting oculomotor orienting and disengagement were completed automatically using gazeAnalysisLib, a library of MATLAB routines for offline analysis of raw gaze data (Leppänen et al., 2015).

#### Oculomotor orienting

Each trial in the oculomotor orienting task started with a 1000-ms blank screen followed by an attention-grabbing animation presented on the center of the screen. The animation was a checkerboard pattern and it rotated into its mirror image after each 250 ms. The pattern remained visible at least 250 ms until the infant looked at it. When the infant looked at the central cue, the cue disappeared and the target (a checkerboard pattern) appeared randomly in one of the four corners of the monitor. The target remained visible 750 ms after the infant’s point of gaze was first recorded in the target area. After that, a picture of a cartoon image or a toy combined by a joyful audio stimulus was presented on the center of the screen as a rewarding stimulus. The eye tracking data was analyzed with the gazeAnalysisLib (Leppänen et al., 2015) to determine the latency of infants’ orienting response (saccadic reaction time) from the central to the target stimulus. Gaze shifts away from the screen and reaction times longer than 1000 ms were excluded.

Infants with 4 or more scorable oculomotor orienting trials were included. The criteria for included trials were: adequate fixation on the central stimulus (i.e., >70 % of the time) during the time preceding a gaze shift, sufficient amount of non-missing data samples (i.e., no gaps longer than 200 ms), and valid information about the time of the gaze transition from the central to the peripheral stimulus (i.e., the transition did not occur during a period of missing data). Trials with gaze shifts within a period that started 150 ms after the onset of the peripheral stimulus and ended 1000 ms after the lateral stimulus onset were used to calculate mean oculomotor orienting latency for each participant (Leppänen et al. 2015).

#### Attention disengagement

Attention disengagement was measured with a gaze-contingent Overlap paradigm, which has been commonly used for measuring infant attention disengagement from various stimuli (e.g., Peltola et al., 2008; 2018). First, a fixation point was presented on the center of the screen until the infant looked at it for 500 ms. After that, a happy or a fearful face, or a non-face stimulus (a phase-scrambled rectangle containing the amplitude and color spectra of the original face stimuli) appeared on the center of the screen for 1000 ms. A distractor stimulus (a vertical checkerboard pattern) was then presented on the left or right side of the screen parallel to the central stimulus for 500 ms. At 250 ms the distractor stimulus changed to its mirror image. After 500 ms, the distractor stimulus disappeared and the central stimulus remained on the screen for 1000 ms, followed by a 1000-ms blank screen before the next trial. The central stimulus conditions (happy or fearful face, or the non-face stimulus) were presented in random order with the restriction that the same stimulus condition was not repeated more than 3 times in a row.

The corneal-reflection eye tracker measured the reaction times for gaze shifts between the central stimulus and the distractor stimulus. A *dwell time index* reflecting attention disengagement was calculated if there were at least 3 scorable trials in each stimulus condition. We calculated the dwell time index using the trials with a gaze shift and trials without a gaze shift, excluding non-scorable trials, and the time interval from the shortest to the longest acceptable saccadic reaction time. Trials with insufficient fixation on the central stimulus (i.e., >70 % of the time) during the time preceding a gaze shift or the end of the analysis period, > 200 ms gaps of valid gaze data, or invalid information about the gaze shift (i.e., with the eye movement occurring during a period of missing gaze data) were excluded from the analyses. For the included trials, attention dwell time on the central stimulus was determined for the period starting 150 ms from the onset of the distractor stimulus and ending 1000 ms after distractor onset. The duration was then converted to a normalized dwell time index score by using the following formula:

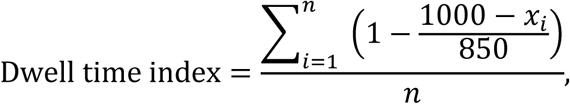

where *x* is the time point of the saccadic eye movement (i.e., last gaze point in the area of the central stimulus preceding a saccade toward the distractor stimulus) and *n* is the number of scorable trials in a given stimulus condition. In this index, the shortest acceptable saccadic reaction time (150 ms) results in 0, and the longest possible saccadic reaction time (or lack of saccade, which is equal to the last measured time point of the central stimulus at 1000 ms) results in a score of 1 (Leppänen et al., 2015). The dwell time index was calculated as the mean of the stimulus conditions (happy or fearful face, or the non-face stimulus) from the accepted trials, and used as the dependent variable in the analyses of attention disengagement. The inclusion criterion was a minimum of 3 acceptable trials in each stimulus condition.

### Eye tracking assessments in Cape Town

Similar to the assessments in Tampere, infants were assessed in a dimly lit room at a private health clinic or a state hospital. A custom MATLAB script (Mathworks, Natick, MA), running on OS X (Apple Inc., CA), and interfacing via Psychtoolbox and Tobii SDK plug-in with the eye tracking system was used for stimulus presentation and data collection (see Pyykkö et al., 2019 for full description of the tests). Eye tracking data were collected by using a Tobii X2-60 or TX60 screen-based eye tracking system (Tobii Technology, Stockholm, Sweden). The procedure for eye tracker calibration was similar for that used in the study in Tampere.

**Oculomotor orienting** was measured with a visual search task (adapted from Kaldy et al.,2011 and described in detail in Pyykkö et al., 2019). In this task, an image of a red apple (5° visual angle) was presented on the center of the screen, accompanied by an *oh* sound. After the infant had looked at the stimulus and 2000 ms had elapsed (or a maximum wait period of 4000 ms had elapsed), the infant saw a blank screen for 500 ms, followed by the re-appearance of the apple in a randomly chosen location. The apple was presented alone (*one-object condition*), among four or eight identical distractors (*multiple-objects condition*), or among four to eight different types of distractors (*conjunction condition*). There was a total of eight trials per condition. To obtain a similar measure for oculomotor orienting as in the Tampere cohort, we extracted orienting latencies from the data collected in the one-object condition, and averaged these latencies for each child to obtain a measure of oculomotor orienting.

In the **disengagement task**, infants saw two stimuli with a 1000 ms onset asynchrony. The first was a picture of a non-face pattern (as above) or a face of a South African female expressing happiness or fear. The ethnicity of the model matched that of the child. The second stimulus was a geometric shape (black and white circles or a checkerboard pattern) laterally on the left or right side of the screen, which was superimposed by a still picture of the first frame of a child-friendly cartoon animation. When the infant shifted gaze to the lateral image or 2000 ms had elapsed, the still picture turned into a dynamic cartoon for 4000 ms. Infants saw a total of 16 trials with non-faces and 16 with faces. As above, the dwell time indices were calculated separately for each of the stimulus conditions by using an automated script.

### Statistical Analyses

We performed the statistical analyses with SPSS version 25.0 (IBM Corp., Armonk, NY). The alpha level for statistical significance was set at .05. All the variables were normally distributed according to Kolmogorov-Smirnov tests. We studied the associations between infant cry and later visual attention with Pearson’s two-tailed correlation coefficients. In the Tampere cohort, we conducted additional analyses to control for the location of the cry sample in the cry bout by including only those infants whose cry sequence was from the beginning of a bout. In the Cape Town cohort, we conducted an additional analysis to control for the possible influence of prenatal SSRI medication by including only those infants without prenatal SSRI exposure. In the Tampere cohort there were no infants with SSRI exposure and in the Cape Town cohort all the cry samples were from the beginning of a cry bout.

## Results

As can be observed from Table 2, the analysis within the Tampere cohort showed a marginal positive correlation between F0 and attention disengagement. When F0 was higher, the dwell time index was larger, i.e., the infants were slower to shift gaze away from the central stimulus to the distractor stimulus. A similar but a larger correlation was observed in the additional analysis including only those infants whose cry sequence was captured from the beginning of a bout. A similar correlation between F0 and attention disengagement was, however, not found in the Cape Town cohort, neither in the primary analyses nor in the analysis where infants with exposure to prenatal SSRI medication were excluded.

**Table 2.** Correlations (Pearson’s *r*) between the F0 parameters and attention outcomes in the two cohorts.

|  |  | Oculomotor orienting | Attention disengagement |
| --- | --- | --- | --- |
| Tampere | F0 | $r = .047, p = .741$ | $r = .288, p = .052$ |
| | F0var | $r = -.102, p = .471$ | $r = -.048, p = .750$ |
| Tampere, beginning of the cry bout | F0 | $r = .249, p = .194$ | $r = .497, p = .008$ |
| | F0var | $r = .115, p = .553$ | $r = -.041, p = .840$ |
| Cape Town | F0 | $r = -.053, p = .726$ | $r = .160, p = .252$ |
| | F0var | $r = -.141, p = .343$ | $r = .038, p = .785$ |
| Cape Town, prenatal SSRI excluded | F0 | $r = -.077, p = .652$ | $r = .141, p = .373$ |
| | F0var | $r = -.073, p = .668$ | $r = -.075, p = .636$ |

There were no statistically significant correlations between F0 and oculomotor orienting in either cohort. Further on, there were no statistically significant correlations between F0 variability and either of the attentional outcome variables. The correlations are presented in Table 2.

## Discussion

The present study is the first to explore the relation of infant cry acoustics to visual attention during the first year. It integrates two previously separate research lines of infant cry analysis and eye tracking based measurement of infant visual attention. Saccadic reaction times during infancy have been suggested to be a basic cognitive skill which mediates the development of more advanced cognitive functions (Dougherty & Haith, 1997). Using these methods, we studied whether higher and more unstable fundamental frequency (F0) of neonatal cry would be connected to visual attention measured with eye tracking in later infancy.

For the most part, the results did not indicate that the F0 of neonatal cry is associated with visual attention at 6 to 8 months of age. The only exception was an association between higher neonatal F0 and slower attention disengagement from central stimuli to peripheral distractor stimuli at 8 months of age in the Tampere cohort. This result was replicated when we controlled for the location of the analyzed cry sample within the cry bout, with a larger correlation observed when the cry was captured from the beginning of the cry bout. The same association was not, however, observed in the Cape Town cohort where the cry was captured at 6 weeks and attention disengagement at 6 months. Furthermore, oculomotor orienting turned out to have no correlation with F0 in either sample. Finally, the variation of F0 did not correlate with the eye tracking outcomes in any of the analyses.

The lack of correlation between neonate cry and oculomotor orienting in the current cohorts could be due to the early maturation of oculomotor orienting, which could result in rather low variability in saccadic reaction times in infants from typically developed cohorts at 6 and 8 months. Indeed, the variability in oculomotor orienting times were moderately low in the current cohorts, and the variability of saccadic reaction times in the attention disengagement task was higher than variability in the oculomotor orienting task. Both F0 and oculomotor orienting have been indicated to reflect later cognitive functions, and the brainstem and vagal nerve to have important roles in their regulation. A distinctive difference between the two attention measurements, i.e., oculomotor orienting and disengagement, is that in the disengagement task, attention needs to be disengaged from a stimulus at fixation, whereas in the oculomotor orienting task, shifts of attention indicate a reaction to a new stimulus appearing on a blank screen. This difference may render the disengagement task cognitively more demanding. Matsuzawa and Shimojo (1997) observed significant differences in the maturation of these two components of attention. Disengagement times decreased markedly from 2 ½ months to 6 months of age, and changed only little from 6 months up to 1 year. In contrast, oculomotor orienting seemed to have matured before the first measurement at 2 ½ months and showed only minor changes within the studied age period. Additionally, saccadic reaction times in the disengagement task at 2 ½ months were approximately twice as long compared to saccadic reaction times in the oculomotor orienting task, and approached the speed of oculomotor orienting at 6 months. Thus, differences in the maturational timing of the two components of attention may have influenced our results, considering that the eye tracking registrations were captured at 6 or 8 months. Future studies may benefit from measuring oculomotor orienting at an earlier age.

Nevertheless, the difference between the maturation of oculomotor orienting and disengagement does not explain why there was a correlation between F0 and attention disengagement only in the Tampere cohort but not in the Cape Town cohort. Considering the lack of correlation between F0 and attention disengagement in the Cape Town cohort we will first pay attention to the age differences at the time of cry recording between the cohorts. Secondly, we discuss the potential influence of early caregiver-infant interaction on the observed pattern of results.

There is a consensus that early cry reflects the development of the central nervous system (LaGasse et al., 2005; Lester, 1987). The regulation of cry goes through prominent changes within the first months of life. Neonatal cry is reflex-like, regulated by the brainstem and not dependent on higher cortical areas, the influence of which increase with age (Newman, 2007). Due to the intense early development of cry regulation, it is thus possible that the variation in cry acoustics at 6 weeks of age does not reflect activity of areas important for visual attention (e.g., brainstem and vagal nerve) to the same extent as cry captured at the neonatal age. Further on, it has been detected that F0 changes as a function of age in normal development. F0 decreases in the first days and weeks, which is usually attributed to the growth of vocal cords. Decrease of F0 is followed by an increase of F0, which is suggested to relate to progressive control of vocalizations due to neurological maturation. However, the exact time course of this process is not clear (Baeck & de Souza, 2007; Gilbert et al., 1996). Consequently, the prominent changes in the regulation of cry as well as normal age-related changes in F0 during the first weeks and months of life highlight the importance for standardizing the age of recording data for cry analysis. For future studies, we suggest recording cry samples at the neonatal period, as this will also make the data comparable to the earlier studies showing associations between neonatal F0 and later cognitive development (Huntington et al., 1990).

Likewise, interaction with the social and physical environment has the potential to influence infants’ cognitive development and cry acoustics. Studies with neonates have shown that the amount and the acoustic qualities of cry may vary as a function of caregiver-infant interaction (Belsky et al., 1984; Lotem & Winkler, 2004), with tactile stimulation and reinforcement learning playing a significant role in the variation of early cry (Cecchini et al., 2007). Consequently, infants learn to adjust their cry to the most effective level according to situational factors. In the current study, the age difference between the cry recordings in Tampere (0-5 days) and Cape Town (6 weeks) may be considered a significant timespan in postnatal development. Additionally, a proportion of Cape Town mothers had psychiatric problems such as depression and anxiety disorders, which are known to influence to mother-child interaction and the development of the child (Field, 2010), and may have further influenced the results. Thus, in future studies, in addition to standardizing the age of cry recordings, measures of early interaction quality are likely important.

The other cry feature that has been associated with later development in previous studies, i.e., variation of fundamental frequency (F0var), was not related to the eye tracking outcomes. A vast amount of studies has observed more varying F0 among preterm and/or low birthweight infants, infants with prenatal risk factors, as well as infants with severe medical or developmental abnormalities. Instead, autism spectrum disorder has been associated with reduced variation in F0. While it is possible that the absence of associations between F0 variation and the outcomes was due to the infants in our study being full term, with normal birth weight, and without any known health or developmental problems, it should be noted that two earlier studies (Huntington et al., 1990; Lester, 1987) detected that more varying F0 was related to lower scores in cognitive outcome measures at several ages even after controlling for the risk factors, including prematurity and prenatal exposure to drugs in the cohorts. As there is currently a little methodological consensus about the determination of F0var, an important line for future studies is to examine the predictive significance of different possibilities in determining F0var, such as index variables or micro-variation measures, e.g., fluctuation or interquartile range.

The influence of the trigger of the cry and the location of the analyzed cry sample in the cry bout need to be considered further. Our additional analysis in the Tampere cohort suggests that the correlation between F0 and attention disengagement is stronger when the cry is captured from the beginning of the cry bout. In a cry triggered by a sudden pain, the F0 is high in the beginning and decreases by time. In contrast, in an inconvenience cry (e.g., hunger, cold, or loneliness) the F0 is lower in the beginning and increases by time if the response to the cry is delayed (Green et al., 1998; Zeskind et al., 1985). In our study the trigger in the Cape Town cohort was either a vaccination or a weight measurement on a scale. In Tampere, the triggers ranged from venipuncture to unknown inconvenience. However, the venipuncture did not trigger a cry in all cases, and there were no other triggers in Tampere which could be considered as painful. In general, attempts to make a taxonomy of different cry triggers face prominent challenges, because evaluating the trigger is more or less subjective except in case of strong pain caused by an external operator. Future studies could benefit from measuring infants’ arousal levels during cry paradigms (e.g., with heart rate or skin conductance measurements) as autonomic arousal is known to influence F0 (Porter et al., 1988; Shinya et al., 2014) and correlate with the experience of pain and inconvenience Stevens et al., 2007). Thus, it would be important to study whether cry samples during increased arousal (like sudden pain, or prolonged cry) are more informative when studying the correlation between cry and later cognitive variables. The representativeness of the used cry samples is important for increasing the accuracy of the used F0 variables. It is possible to use several recordings or multiple samples within a recording. However, also in this approach the influence of arousal and the location of the used sample in the whole cry sequence should be taken into consideration.

The reliability of the correlation estimates in the current study was limited by the number of infants included in the analyses, and it will be important to replicate the findings with a larger sample. Additionally, the tasks for measuring oculomotor orienting and attention disengagement were not completely identical in Tampere and Cape Town, which may have influenced the results. Notwithstanding these limitations, a clear strength of the study is the integration of two previously separate research lines of infant cry analysis and eye tracking based measurement of infant visual attention. Both methods enable registrations at a very early age which is a prominent advantage in evaluating early cognitive development. Methodological development (e.g., Naithani et al., 2018) has also enabled reliable segmentation and acoustic analyses of cry sounds captured in various settings, broadening the possibilities of neonatal cry analysis.

Monitoring early cognitive development is an important, yet challenging part of pediatric practice, which increases the need for applicable methods that can be administered at an earlier age than more elaborate psychological assessments. Research on the potential of using infant cry to predict cognitive development is still scarce and often conducted with small samples. In the present study with typically developing infants, the results suggested a small association between neonatal cry F0 and later visual attention disengagement. Whether variations in infant cry characteristics are sufficiently distinct in typically developing populations to be used as a supporting tool for monitoring cognitive skills requires further research. Nevertheless, detecting normative variation in infant cognition is vitally important especially for identifying children with milder forms of atypical cognitive development. To reach this population for appropriate support, rehabilitation and health care as early as possible would be especially important.

## Notes

### Competing Interest Statement

The authors have declared no competing interest.

